# The crystal structure of the Shethna Protein II (FeSII) from *Azotobacter vinelandii* suggests a domain swap

**DOI:** 10.1101/2024.01.30.575082

**Authors:** Burak V. Kabasakal, Ciaran R. Mcfarlane, Charles A. R. Cotton, Anna Schmidt, Andrea Kung, Lucas Lieber, James W. Murray

**Affiliations:** Department of Life Sciences, Imperial College, London, London SW7 2AZ, United Kingdom; Turkish Accelerator and Radiation Laboratory, Gőlbaşı, 06830, Ankara, Turkiye; Cambrium GmbH, Max-Urich-Strasse 3, 13355, Berlin, Germany; Bioheuris Inc. 1100 Corporate Square Dr, St. Louis, MO 63132 USA

**Keywords:** nitrogen fixation, oxygen protection, structural biology, domain swap

## Abstract

The *Azotobacter vinelandii* FeSII protein forms an oxygen resistant complex with the nitrogenase MoFe and Fe proteins. FeSII is an adrenodoxin type ferredoxin, forming a dimer in solution. Previously, the crystal structure was solved (Schlesier et al., 2016), with 5 copies in the asymmetric unit. One copy is a normal adrenodoxin domain, forming a dimer with its crystallographic symmetry mate. The other four copies are in an “open” conformation with a loop flipped out exposing the 2Fe-2S cluster. The open and closed conformations were interpreted as oxidised and reduced, and the large conformational change in the open configuration allowed binding to nitrogenase. We have independently solved the structure of FeSII in the same crystal form. Our positioning of the atoms in the unit cell is similar to the earlier report. However, our interpretation of the structure is different. We interpret the “open” conformation as the product of a crystallization-induced domain swap. The 2Fe-2S cluster is not exposed to solvent, but in the crystal its interacting helix is replaced by the same helix residues from a crystal symmetry mate. The domain swap is complicated, as it is unusual in being in the middle of the protein rather than at a terminus, and it creates arrangements of molecules that can be interpreted in multiple ways. We caution that crystal structures should be interpreted in terms of the contents of the entire crystal rather than one asymmetric unit.

## Introduction

Nitrogen is an essential component of DNA and proteins. Nitrogen gas makes up 80% of the atmosphere, but it is very unreactive. Fixing this unreactive nitrogen is an essential biological and industrial process. Biological nitrogen fixation is catalysed by the enzyme nitrogenase, which reduces nitrogen gas to ammonia. The nitrogen reduction is catalysed by the MoFe protein, a (NifDNifK)_2_ tetramer, containing molybdenum. The MoFe protein is in turn reduced by the dinitrogenase reductase, the Fe protein, a dimer of NifH. MoFe has two oxygen sensitive metalloclusters, the P-cluster and the FeMoCo, containing the molybdenum atom. The Fe protein has a low potential 4Fe-4S cluster at the surface bridging the dimer interface. Both nitrogenase components are irreversibly inhibited by oxygen, which reacts with the low potential iron-sulfur clusters. MoFe has a half life of around 10 minutes in air, and the Fe protein has a half life of around 45 seconds (Robson, 1979; Robson and Postgate, 1980). The MoFe:Fe protein complex is partially protected from oxygen, relative to each protein individually (Schlesier et al., 2016). As well as direct inactivation of the nitrogenase, oxygen reactions with nitrogenase proteins probably generate damaging reactive oxygen species (Maier and Moshiri, 2000).

Organisms that fix nitrogen, known as diazotrophs, use several strategies to protect themselves from oxygen. Some are obligate anaerobes or micro-aerobes. Some diazotrophic aerobes such as *Azotobacter* species use “respiratory protection”, a fast respiratory rate to consume oxygen. A second mechanism is “conformational protection”. It was noticed that a nitrogen fixing culture of *Azotobacter chroococcum* would stop fixing oxygenated by strong agitation, but when the agitation stopped, the culture would resume nitrogen fixation at the same rate as before (Hill et al., 1972). This result implied that the nitrogenase was being temporarily inactivated, by a “conformational” mechanism.

In the 1960s, Shethna and colleagues isolated several non-heme iron proteins from *Azotobacter vinelandii* (Shethna et al., 1968). The second of these proteins became known as the Shethna protein, or FeSII. Early work on purifying nitrogenase found a pink protein that associated with nitrogenase, and provided some oxygen protection (Kelly et al., 1967). This pink protein was then identified as the FeSII protein (Bulen and LeComte, 1972; Haaker and Veeger, 1977). FeSII is the basis of the conformational protection (Robson, 1979). It forms a ternary complex with the MoFe and Fe proteins that is catalytically inactive, but from which active nitrogenase can be recovered, with a half life in air of about an hour (Robson, 1979). FeSII is not essential for *A. vinelandii* viability, even under nitrogen fixing conditions, but the nitrogenase was more sensitive to inactivation in an FeSII knockout mutant (Moshiri et al., 1994).

FeSII has only been shown to have protective activity in *A. vinelandii and A. chroococcum*, although FeSII-like functionality has been proposed for other ferredoxins associated with nitrogenase operons, for example FdxD in *Rhodobacter capsulatus* (Hoffmann et al., 2014) and a *Guconacetobacter* ferredoxin (Lery et al., 2010; Ureta and Nordlund, 2002). It is possible that protective function has appeared more than once, or that similar mechanisms exist in other diazotrophs, but have not yet been identified.

FeSII is a 13 kDa adrenodoxin type ferredoxin protein with a 2Fe-2S cluster, forming a dimer in solution, the midpoint potential is -262 mV (Moshiri et al., 1995). The crystallization was published in 1995 (Moshiri et al., 1995), in an orthorhombic crystal form, but no structure was solved. Recently Schlesier *et al* (Schlesier et al., 2016) reported a 2.2 Å crystal structure of FeSII in the same orthorhombic crystal form. The observed FeSII crystal form has five copies in the asymmetric unit. One of them is in a “closed conformation”, forming a “closed” dimer with a crystallographic symmetry mate. The closed conformation is similar other adrenodoxin proteins, such as the monomeric *E. coli* Fdx (Kakuta et al., 2001). The other four copies in the asymmetric unit are in an “open” from. In the open form, the so-called N-loop from residues 59-96 is flipped out from the structure so that the 2Fe-2S cluster is revealed. Specifically, the “lid helix” from residues 68-78 is moved away from the cluster. The four open FeSII were interpreted as two dimers of the same type as the closed form, but with the N-loop flipped out. Schlesier et al proposed that this conformational change is triggered by oxidation of the cluster, and the change enables FeSII to form the protective ternary complex with MoFe and Fe proteins. The open form of FeSII has not been observed in crystals of any other adrenodoxin type domains.

Protein domain swapping is when two protein molecules exchange equivalent regions in their structures (Liu and Eisenberg, 2002; Wodak et al., 2015). In the simplest form, two adjacent monomers swap a region to form a domain-swapped dimer. Some proteins have evolved natural domain swaps, relative to an unswapped ancestor (Cahyono et al., 2020). In other cases the swap is functional, or may be involved in disease (Rousseau et al., 2012). Domain swapping may also be induced by chemical treatment (Cahyono et al., 2020) or crystallization. For example in a crystallised PAS domain, one of the six non-crystallographic symmetry copies had a domain-swapped C-terminal alpha-helix (Emami et al., 2009). In cyanobacterial Psb29 (Bečková et al., 2017), only one of three crystal forms had a domain-swapped N-terminal alpha-helix (Bečková et al., 2017). In these cases, the domain swapped structure may be regarded as a crystallization artefact, and the swapped structure is not biologically relevant. In most domain-swapped structures the swap is of one end of the structure containing the N- and C-terminus. Swaps of internal regions are rarer, but have been observed, such as in the structure of IX/X-binding protein from snake venom (Mizuno et al., 1997), which has a loop exchange heterodimer, relative to a the homologous rat mannose binding protein (Weis et al., 1991).

We have independently solved the structure of the FeSII protein in the same crystal form as previously, to higher resolution. Our atoms are in similar positions to the earlier structure, but our interpretation of the open form of the protein is very different. We believe that the open form is caused by a crystallization-induced domain swap. Here, we present evidence that the “open” conformation of FeSII in the crystal structure is best interpreted as a domain-swapped structure.

## Results

### FeSII structure

We independently solved the FeSII structure using Fe-SAD to a final resolution of 1.65 Å in the same orthorhombic P2_1_2_1_2 crystal form as observed in 1995 (Moshiri et al., 1995) and 2016 (Schlesier et al., 2016). Data and refinement information are in Table S1. The asymmetric unit contains five copies of FeSII. One is in the closed conformation similar to other adrenodoxins. The other four are in the open conformation. Although the crystal form is the same, we have placed the open protein chains in different symmetry positions to highlight our interpretation of the structure. The structures are similar, with an RMSD of 1.1 Å over the closed chain between this work and Schlesier et al (2016), and 0.4 Å between the best matching open chains. The closed chain is more complete than before, with a gap from residues 82-92 in the N-loop. In the open chains, the N-loop is ordered, and the entire chain is visible, apart from the N- and C-termini.

### Interpretation of crystal structure

In this crystal form the “closed” chain E forms a dimer with its crystallographic symmetry mate E’ (-X+1, -Y, Z). All four open chains form the “open dimer”. Chains A and B form this dimer. Chain C forms this dimer with Chain D from the X,Y,Z-1 symmetry mate (D’). Chains C and D form another extended interface forming we call the “domain-swap” dimer. Chain A also forms this domain swap dimer with the B chain related by X,Y,Z-1 (B’). All open chains are involved in both the “open” and “domain-swap” dimers. Fig 1 shows the dimers in the crystal and the chains forming them for two unit cells.

**Figure 1:**
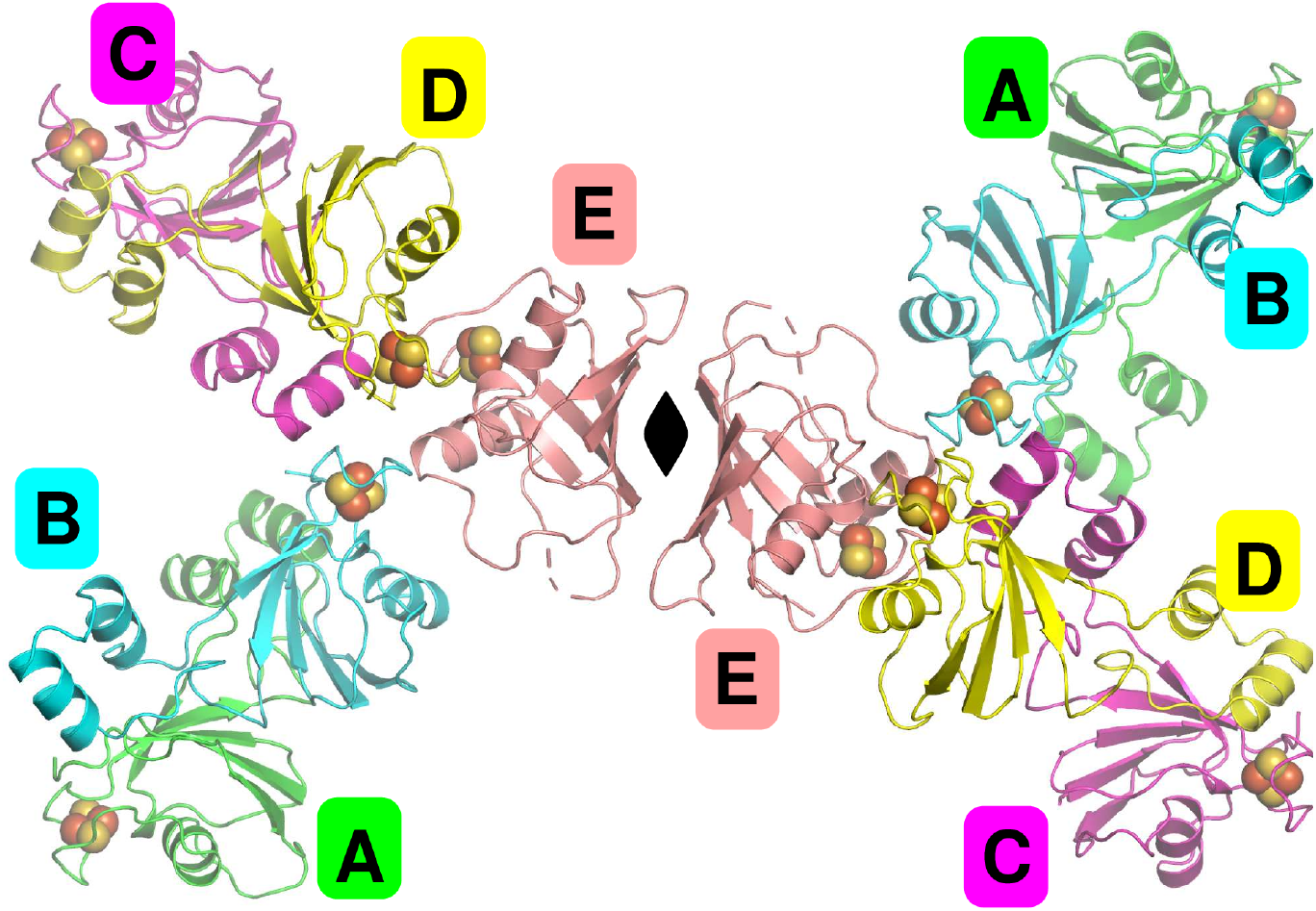
The contents of two unit cells of the FeSII protein, showing the five unique chains A, B, C, D.

The work by Schlesier et al proposed that the “lid helix”, moves away from the 2Fe-2S cluster in the open dimer, exposing it to solvent. However, in the crystal, in each open chain the lid helix is replaced by a lid helix from a symmetry related chain. Fig. 2a shows the lid helix in place for chain E, the closed chain. For the open chain C, Fig. 2b shows the lid helix in an equivalent position, but coming from the domain-swap dimer partner chain D in the crystal. In chains A and B, the lid helix is similarly replaced by symmetry related chains B’ and A’ respectively.

**Figure 2:**
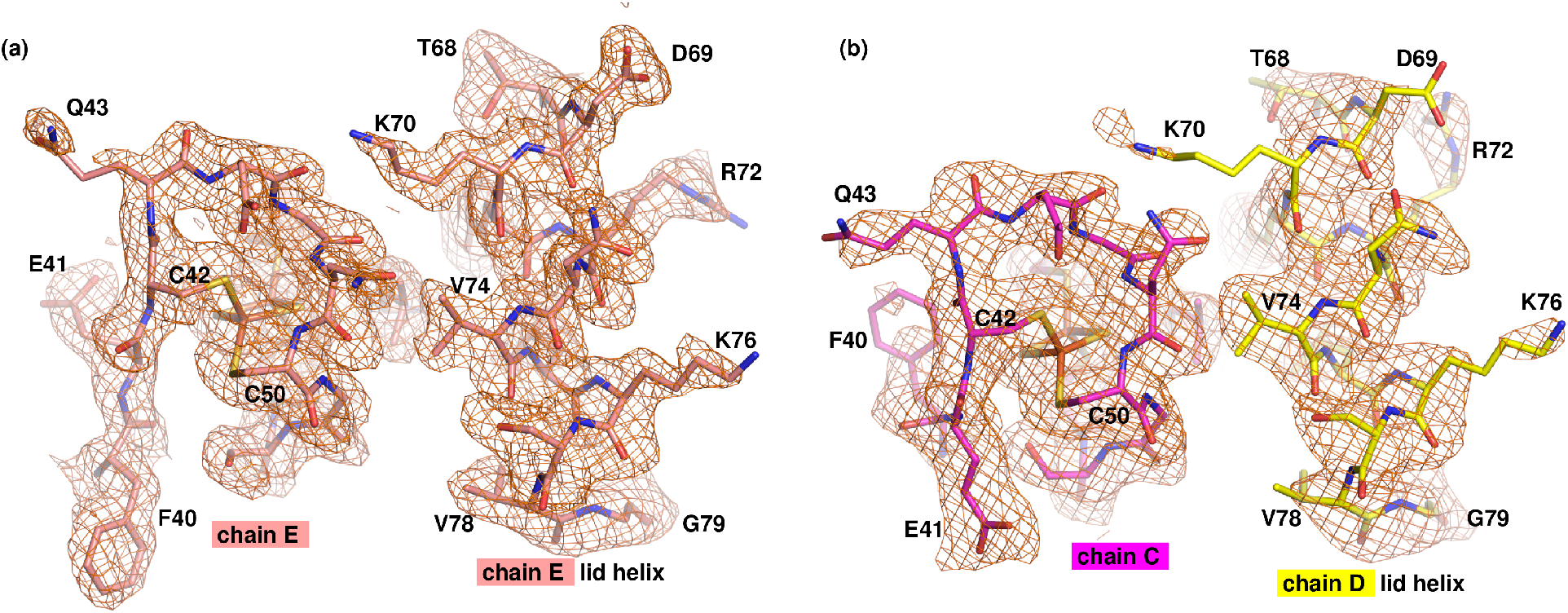
Region around the 2Fe-2S cluster for closed and open chains. For chain E (salmon) in the crystal, the lid helix from chain E covers the iron-sulfur cluster. For chain C (magenta), the lid helix is flipped out in the open conformation, but the cluster is still capped by the lid helix from chain D. Weighted 2F_o_-F_c_ density is contoured at 0.9σ and selected residues are labelled.]

The predicted interface areas, and binding energies for the dimers that exist in the crystal are shown in Table 1. The calculated area of the closed dimer interface is 490 Å^2^.

**Table 1:** Interface areas in Å^2^, calculated as half the difference in total accessible surface areas of isolated and interfacing structures.

| Interface | Chains | Interface Area / Å<sup>2</sup> | Predicted binding energy / kcal mol<sup>-1</sup> |
| --- | --- | --- | --- |
| OPEN 1 | A:B | 1170 | -9.3 |
| OPEN 2 | C:D' [X,Y,Z-1] | 1080 | -8.4 |
| MEAN OPEN | A:B, C:D' | 1120 | -8.9 |
| DOMAIN SWAP 1 | C:D | 983 | -16.6 |
| DOMAIN SWAP 2 | A:B' [X, Y, Z-1] | 960 | -17.0 |
| MEAN DOMAIN SWAP | C:D, A:B' | 970 | -16.8 |
| CLOSED | E:E' [-X+1, -Y,Z] | 490 | -4.8 |

In the open conformation, the “lid helix” (residues 68-79), as part of the longer “N-loop” (residues 59-96) is folded away from the central domain. It forms additional contacts with the closed dimer partner around Pro12, expanding the closed dimer interface. The calculated area of the closed dimer interface is 490 Å^2^, but the mean open dimer interface area is 1120 Å^2^. In the crystal, the open dimers are also involved in the domain swap dimer. This dimer is interface is primarily made up of residues of the opened out N-loop, but has a mean area of 970 Å^2^, and is the most energetically favoured of the interfaces. These predicted energies suggest that the formation of the open and domain swap interfaces is energetically favoured in the crystal.

The dimers in the FeSII crystal are shown in Fig. 3 in cartoon and schematic view.The replacement of the lid helix by the lid helix from a symmetry mate (red in Fig 3), may be interpreted as a domain swap of the N-loop. This domain swap is unusual in that it involves an internal region of the protein rather than an N-or C-terminus. The swap is also unusual in that it creates an extensive new dimer interface, in addition to the dimer interface that is present in the open and closed dimers. Given that these two interfaces can only occur together in the crystal, we suggest that this open conformation of FeSII is crystallization induced. The flexibility of the “hinge’’ regions proposed earlier may allow this protein to change shape under the constraints of crystal lattice formation to a more energetically favoured conformation that may not be biologically relevant.

**Figure 3:**
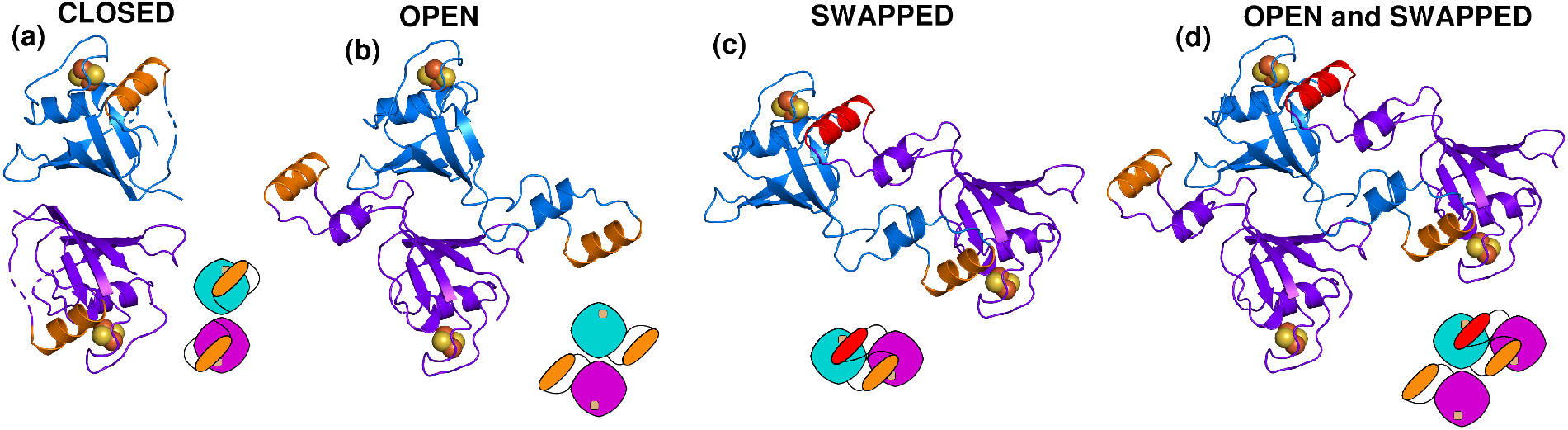
Cartoon and schematic view of the dimers of FeSII in the P21212 FeSII crystal form. (a) closed dimer (chains E and E’), (b) open dimer (chains A and B) (c) domain swap dimer (chains A and B’). (d) Open and domain swap dimers (A, B and B’) The chains are shown in blue and purple, with the lid helix (residues 68-79) in orange, except for the top lid helix when interpreted as part of the domain swap dimer, when it is shown in red.

Schlesier et al propose that the formation of the open dimers is induced by the oxidation of the 2Fe-2S cluster. Our protein and crystals were prepared aerobically, and our protein has an absorption spectrum corresponding to oxidised protein (Fig S1) so even the closed chain should be oxidised, and so any conformational change is not driven by differential oxidation states of the 2Fe-2S cluster.

## Discussion

Previous research has claimed that the FeSII protein adopts a radically changed conformation in the oxidised state, based on the crystal structure and small shifts in retention time in size-exclusion chromatography. We have independently solved the FeSII crystal structure at 1.65 Å resolution. Based on examination of the full unit cell of the crystal, we conclude that the “open’’ structure seen in the crystal is most likely due to crystallization-induced domain-swapping. Previous work on the homologous ferredoxin VI from *Rhodobacter capsulatus* (Sainz et al., 2006) solved structures of the oxidised and reduced protein (Armengaud et al., 2001; Sainz et al., 2006) and only small conformational changes were seen, mainly around the iron-sulfur cluster. It is likely that the interaction of FeSII with nitrogenase is also governed by only small conformational changes. We await structure determination of the full MoFe:Fe:FeSII protein complex for confirmation.

## Materials and Methods

### Overexpression and Purification of FeSII

A synthetic gene coding for the FeSII from *Azotobacter vinelandii* (AVIN_39700) was synthesised by DNA 2.0 (California, USA), and sub-cloned into a pRSET-A vector (Invitrogen, Carlsbad, CA, USA) with no tag. The FeSII gene was PCR amplified with primers FESII_FOR 5’-TATTA<u>CATATG</u>GCGACGATCTATTTCAGCAGC-3’ and FESII_REV 5’-ATTAT<u>CTCGAG</u>TTATCAGGCGCCACCCG-3’ (restriction sites underlined) and the product digested with NdeI and XhoI, and then ligated into vector cut with NdeI and XhoI restriction enzymes.

FeSII protein was produced by transformation of the vector into chemically competent *E. coli* KRX cells (Promega, Wisconsin, USA), and cultivation in 1 L Terrific Broth medium supplemented with 100 μg/l ampicillin. Expression was induced at an OD600 of 0.6-0.8 with 1 g/l L-rhamnose monohydrate. After overnight induction at 18° C, the cells were harvested by centrifugation, and resuspended in 10 mM HEPES pH 7.4, 1 mM MgCl_2_. Cell disruption was performed by sonication. FeSII was purified with a modification of a published method (Moshiri et al., 1995). 0.4 % (w/v) NaCl and 11% (w/v) PEG 8000 were added as powder to the lysate. They were mixed gently for 45 min at 4 °C, and the cell debris was separated by centrifugation. The red supernatant was filtered through a 0.2 μm filter, and purified by anion exchange chromatography on a Q Sepharose column equilibrated against 10 mM HEPES, pH 7.4, 1 mM MgCl_2_. FeSII was eluted by using a linear gradient of MgCl_2_ from 0 mM to 50 mM over 10 column volumes. The fractions corresponding to FeSII protein were pooled and concentrated, and further purified by size-exclusion chromatography (HiLoad 16/600 Superdex 200 PG column) in 10 mM HEPES, pH 7.4, 5 mM MgCl2, 100 mM NaCl.

### Structure Determination of FeSII

FeSII was crystallized using hanging drop vapor diffusion with an equal mix of 0.1 M HEPES pH 8.0, 0.2 M sodium citrate, 24%, (w/v) PEG3350, 2% (v/v) 1-butanol and 40 mg/ml FeSII. Crystals were cryoprotected for 30 s in the mother liquor solution with 30% volume added PEG400, and flash cooled in liquid nitrogen. X-ray diffraction data were collected at 100 K on beamline I03 at Diamond Light Source, UK. For phase determination, a data set was collected at the iron K-edge at a wavelength of 1.734 Å. The data were processed and scaled with xia2 (Winter, 2010) and XDS (Kabsch, 2010). The structure was phased with Fe-SAD using the intrinsic Fe atoms of the protein using AutoSol of the phenix package. Phases were improved with density modification in DM (Cowtan, 2010), and a preliminary model built with Buccaneer (Cowtan, 2006). The model was rebuilt manually in Coot (Emsley et al., 2010), cycling with refinement in phenix.refine (Adams et al., 2010). This model was then refined and rebuilt against a higher resolution native dataset at 1.65 Å. The structure was validated with MolProbity (Williams et al., 2018). Final data collection and refinement statistics are given in Supporting Table 1.

Dimer surface areas and interaction energies were predicted with PISA (Krissinel, 2010; Krissinel and Henrick, 2007). Molecular figures were produced in PyMol (Delano, 2002).

## Acknowledgements

The authors would like to thank Diamond Light Source for beam time (proposal mx12579), and the staff of beam lines I02, I03 and I24 for assistance with crystal testing and data collection. This work was funded in part by the BBSRC/NSF Nitrogen Ideas Lab: Oxygen-Tolerant Nitrogenase (BB/L011468/1). The crystallization facility at Imperial College was funded by BBSRC (BB/D524840/1) and the Wellcome Trust (202926/Z/16/Z). CRM was funded by a BBSRC Doctoral Training Programme (BB/J014575/1), as was CARC (BB/F017324/1).

## Author contributions

JWM, LL, CARC, AK, AS, CRM, BVK, performed research. JWM drafted the paper in consultation with the other authors.

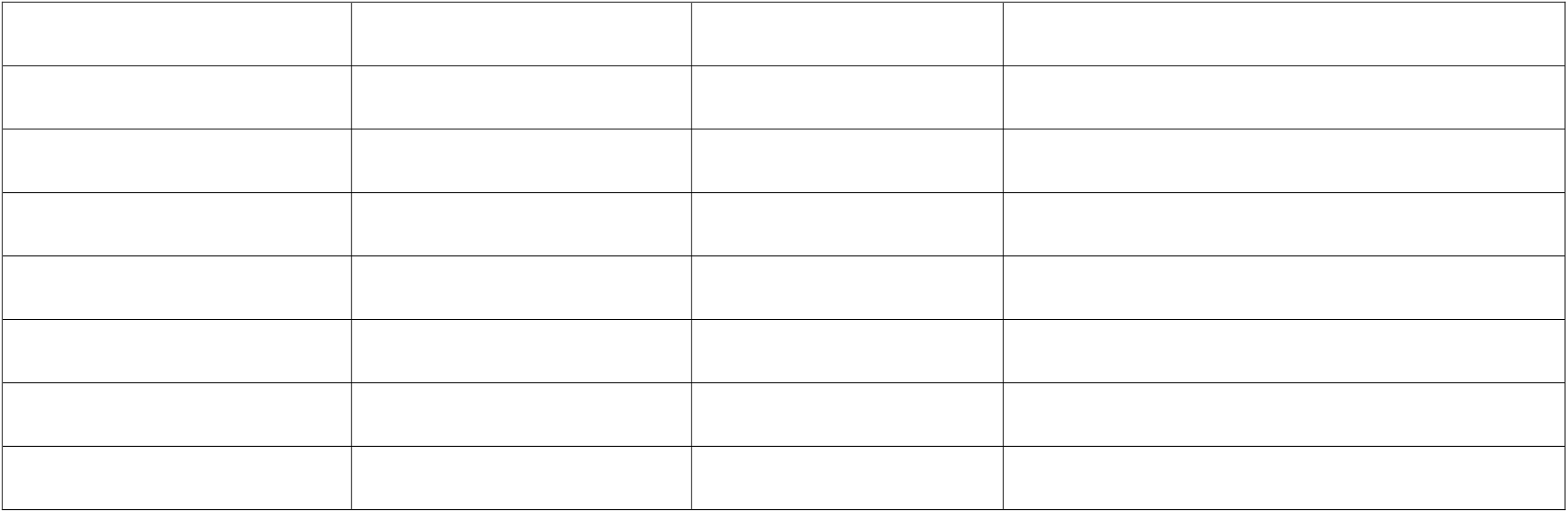

## SUPPLEMENTARY INFORMATION

**Table S1.**
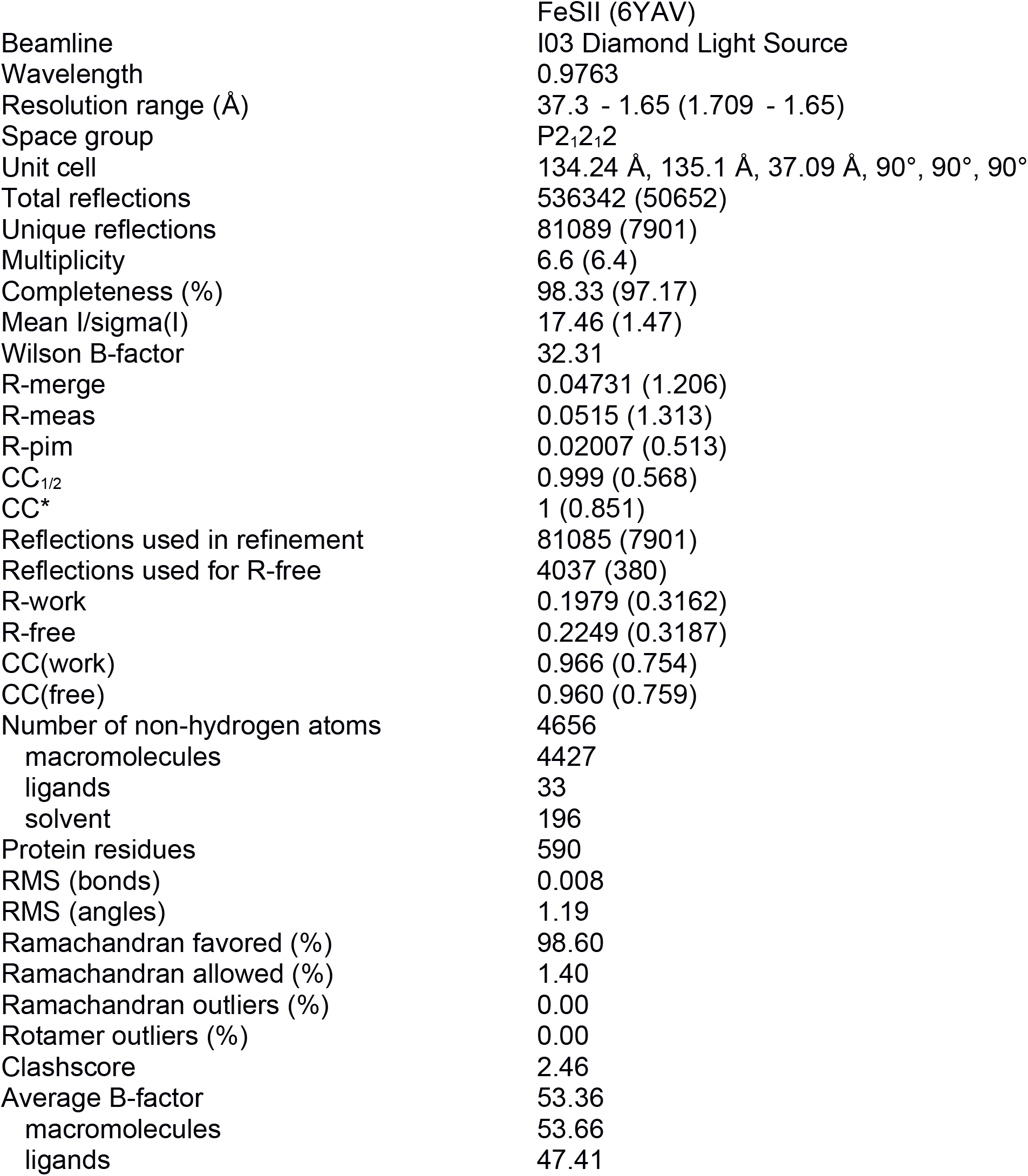
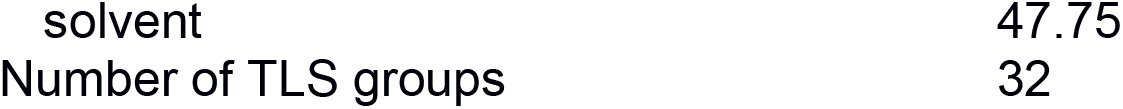
FeSII Data collection and refinement statistics. Statistics for the highest-resolution shell are shown in parentheses.

**Figure S1.**
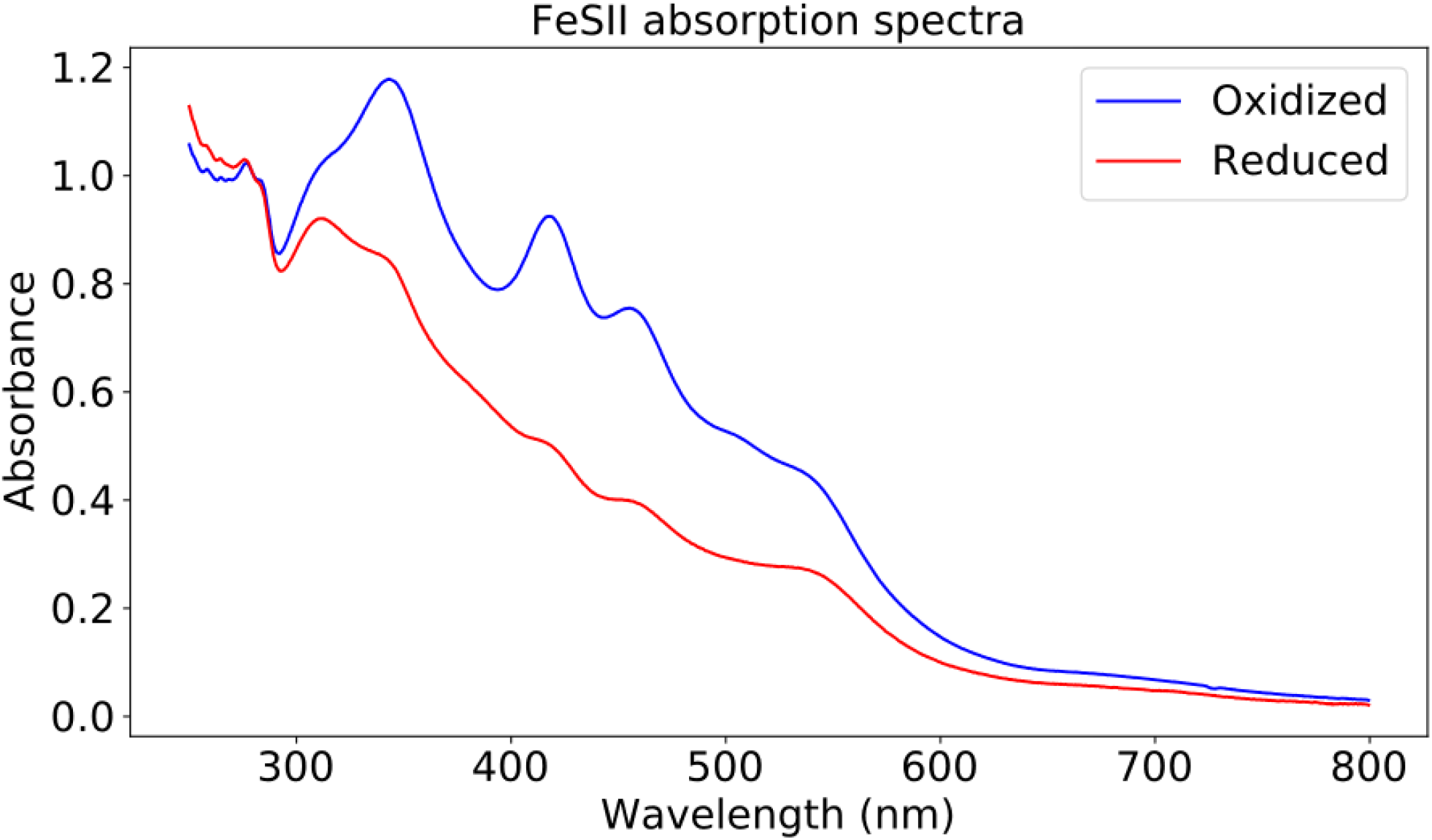
UV-VIS Absorption spectrum of FeSII (1.35 mg/ml) oxidised (blue) and reduced (red) 100 mM HEPES pHh 7.4, 5 mM MgCl2, 100 mM NaCl pH 7.4. Oxidised protein was FeSII prepared in air, reduced protein was prepared by treating anaerobically with 100 μM mM final concentrationsodium dithionite, with excess dithionite removed by repeated centrifugal concentration with a 10 kDa cutoff membrane.

